# Sympathetic structural and electrophysiological remodeling in a rabbit model of reperfused myocardial infarction

**DOI:** 10.1101/2024.06.05.597494

**Authors:** Amanda Guevara, Charlotte E.R. Smith, Lianguo Wang, Jessica L. Caldwell, Srinivas Tapa, Samantha D. Francis Stuart, Betty W. Ma, G. Andre Ng, Beth A. Habecker, Zhen Wang, Crystal M. Ripplinger

**Affiliations:** Department of Pharmacology, University of California Davis, Davis, CA, USA; Campus Veterinary Services, University of California Davis, Davis, CA, USA; Department of Cardiovascular Sciences, University of Leicester, Leicester, UK; National Institute for Health & Care Research, Leicester Biomedical Research Centre, Leicester, UK; Glenfield Hospital, Leicester, UK; Department of Chemical Physiology and Biochemistry, Oregon Health and Science University, Portland, OR, USA; Department of Medicine and Knight Cardiovascular Institute, Oregon Health and Science University, Portland, OR, USA; Shantou University Medical College, Shantou, China

**Keywords:** Arrhythmia, chondroitin sulfate proteoglycans, hypo-innervation, myocardial infarction, sympathetic nerve stimulation

## Abstract

Chondroitin sulfate proteoglycans (CSPGs) inhibit sympathetic reinnervation in rodent hearts post myocardial infarction (MI), causing regional hypo-innervation that is associated with supersensitivity of β-adrenergic receptors and increased arrhythmia susceptibility. To investigate the role of CSPGs and hypo-innervation in the heart of larger mammals, we used a rabbit model of reperfused MI and tested electrophysiological responses to sympathetic nerve stimulation (SNS). Innervated hearts from MI and sham rabbits were optically mapped using voltage and Ca^2+^-sensitive dyes. SNS was performed with electrical stimulation of the spinal cord and β-adrenergic responsiveness was tested using isoproterenol. Sympathetic nerve density and CSPG expression were evaluated using immunohistochemistry. CSPGs were robustly expressed in the infarct and border zone of all MI hearts, and the presence of CSPGs was associated with reduced sympathetic nerve density in the infarct vs. remote region. Action potential duration (APD) dispersion and susceptibility to ventricular tachycardia/fibrillation (VT/VF) were increased with SNS in MI hearts but not in sham. SNS decreased APD_80_ in MI but not sham hearts, while isoproterenol decreased APD_80_ in both groups. Isoproterenol also shortened Ca^2+^ transient duration (CaTD_80_) in both groups but to a greater extent in MI hearts. Our data suggest sympathetic remodeling post-MI is similar between species, with CSPGs associated with sympathetic hypo-innervation. Despite a reduction in sympathetic nerve density, the infarct region of MI hearts remained responsive to both physiological SNS and isoproterenol, potentially through preserved or elevated β-adrenergic responsiveness, which may underly increased APD dispersion and susceptibility for VT/VF.

**NEW & NOTEWORTHY:** Here we show that CSPGs are present in the infarcts of rabbit hearts with reperfused MI, where they are associated with reduced sympathetic nerve density. Despite hypo-innervation, sympathetic responsiveness is maintained or enhanced in MI rabbit hearts, which also demonstrate increased APD dispersion and tendency for arrhythmias following sympathetic modulation. Together, this study indicates that the mechanisms of sympathetic remodeling post-MI are similar between species, with hypo-innervation likely associated with enhanced β-adrenergic sensitivity.

## INTRODUCTION

Myocardial infarction (MI) is a prevailing cause of cardiovascular disease and is associated with substantial risk of ventricular arrhythmias (1, 2). In addition to myocyte loss due to ischemia following MI, there is also significant remodeling of cardiac sympathetic innervation. Post-MI sympathetic hypo-innervation is associated with worse arrhythmia outcomes and sudden cardiac death (3-5), with experimental studies indicating that non-uniform sympathetic nerve density along with changes in adrenergic sensitivity may be arrhythmogenic (6-8).

Electro-anatomical mapping of hearts from human MI patients has shown areas of sympathetic hypo-innervation are associated with β-adrenergic supersensitivity and increased dispersion of the activation recovery interval, a surrogate for action potential duration (APD), during sympathetic stimulation (9). We have similarly shown β-adrenergic supersensitivity, Ca^2+^ mishandling, increased APD dispersion, and increased arrythmia susceptibility in hypo-innervated post-MI mouse hearts (7), along with changes in β-adrenergic and electrophysiological responses in areas of sympathetic hypo-innervation even in the absence of ischemia or MI (10).

Previous work has suggested a role for chondroitin sulfate proteoglycans (CSPGs) in modulating cardiac innervation. In mouse models of reperfused MI, CSPGs present in the infarct inhibited sympathetic nerve reinnervation in the infarct and the adjacent myocardium (7, 11). However, mouse heart size, electrophysiology, and arrhythmia patterns differ from that of larger species (12, 13), including human, where the role of CSPG-associated hypo-innervation post-MI has yet to be explored. To address this, we used a translationally-relevant rabbit model that has similar electrophysiology and arrhythmogenesis to humans (14-17) to investigate electrophysiological responses to sympathetic stimulation and determine if CSPGs and associated sympathetic hypo-innervation are present in a larger animal model of reperfused MI.

## MATERIALS AND METHODS

### Ethical Approval

All procedures involving animals were approved by the Animal Care and Use Committee of the University of California, Davis (protocol #20991), and adhered to the Guide for the Care and Use of Laboratory Animals published by the National Institutes of Health (NIH publication no. 85-23, revised 2011).

### Rabbit MI model

New Zealand White rabbits (4–12 months old, 2.6–3.0 kg, N=22; Charles River Laboratories) were housed for ≥1 month before the study with *ad libitum* access to food and water. Rabbits were randomly assigned to sham or MI surgery at a ratio of ∼1:3 to account for MI-associated mortality. For MI surgery, rabbits were sedated with butorphanol and/or acepromazine. Anesthesia was induced with ketamine and diazepam intravenously (IV) via an ear vein catheter and maintained with inhaled isoflurane (∼2%) via a ventilator for the duration of surgery. A ∼4 cm incision was made at the 4^th^ intercostal space and a chest retractor was placed to expose the heart and lungs. The pericardium was carefully separated to visualize the left coronary artery and the descending branch of the left circumflex artery was ligated with a suture midway between the apex and base for 45–60 min (18, 19). During the ligation, the retractor was removed, and the chest was closed. A bolus of lidocaine (0.5–1 mg/kg) was IV infused before ligation to prevent ventricular fibrillation (VF). In the event of VF, rabbits were given an additional bolus of lidocaine (0.5 mg/kg, IV) and hearts were manually massaged by hand for recovery. Sham surgeries were performed by passing the suture under the coronary artery without ligation. Heart rate (HR), body temperature and blood pressure were monitored during surgery, with body temperature maintained at 37°C by a temperature-controlled warming pad. Ringer’s lactate was given via IV infusion during surgery to ensure adequate hydration. Once surgery was completed, rabbits were returned to housing and monitored. Enrofloxacin (5 mg/kg) was given once to prevent infection, with buprenorphine (0.05–0.1 mg/kg) given immediately post-surgery and then twice daily for 48 hours and Meloxicam given once daily for 5 days. Animals remained in housing until experimental use 10±1 days post-surgery.

### Sample Sizes

Of the 17 MI operated rabbits, 10 (7 males, 3 females) survived surgery, with the 7 deaths associated with ventricular arrhythmias during reperfusion despite lidocaine administration and defibrillation. Although more male than female animals survived, there was no sex difference in survival rate (*P=*0.1534). Of the 10 surviving animals with MIs, 8 hearts were successfully perfused for mapping. However, due to technical limitations of the innervated heart preparation, visualization the infarct region within the mapping field of view and performance of all measurements or pacing protocols was not possible in every heart. As such, the number of animals is variable for some parameters and detailed within each figure legend.

### Innervated Heart Preparation

Innervated hearts were prepared as previously described (20-22). Briefly, on day 10±1 post-surgery, rabbits were administered heparin (1000 IU), then an overdose of pentobarbital sodium (>100 mg/kg) IV via an ear vein catheter. Upon deep anesthesia, the front of the ribcage was removed, and the pericardium gently pulled apart and away from the heart, and the chest filled with ice-cold cardioplegia (in mmol/l: NaCl 110, CaCl_2_ 1.2, KCl 16, MgCl_2_ 16, and NaHCO_3_ 10). The descending aorta was then cannulated with an 8-gauge cannula and flushed with ice-cold cardioplegia. The heart and posterior rib cage were removed with the thoracic spinal column intact by cutting the cervical spine at C1 and the thoracic spine at T12. The preparation was then dissected from surrounding tissues and submerged into ice-cold cardioplegia. The innervated heart was moved to a glass-jacketed chamber and perfused via descending aorta with Tyrode’s solution at 37 °C: (in mmol/L: NaCl 128.2, CaCl_2_ 1.3, KCl 4.7, MgCl_2_ 1.05, NaH_2_PO_4_ 1.19, NaHCO_3_ 20 and glucose 11.1). The excitation-contraction uncoupler blebbistatin (Tocris Bioscience; 10–20 µM) (23) and the nondepolarizing skeletal muscle paralytic vecuronium bromide (Cayman Chemical; 5 µM) were added to the perfusate to eliminate motion artifacts during optical recording. Perfusion pressure was maintained at 40-60 mmHg by adjusting the flow rate (∼100-120 ml/min). Three Ag/AgCl needle electrodes were positioned in the bath (two in the thoracic cavity and one grounded in the chamber) for continuous ECG recording.

### Dual Optical Mapping

Dual optical mapping of intracellular Ca^2+^ and transmembrane potential (V_m_) was performed as previously described (13, 20, 22, 24). Briefly, hearts were loaded with 1 mg/ml Rhod-2 AM in 0.5 mL of DMSO containing 10% pluronic acid (Molecular Probes) followed by 50 µL 1 mg/ml RH237 in DMSO (Molecular Probes). The dyes were excited at 531±20 nm (LEX-2, SciMedia). Emitted fluorescence was collected using a THT-macroscope (SciMedia). Emission signals were split with a dichroic at 630 nm, longpass filtered at 700 nm (V_m_) and bandpass filtered at 590±16 nm (Ca^2+^). Images were acquired at 1 kHz with a 31×31 mm (100×100 pixels) field of view using CMOS cameras (MiCam Ultima-L, SciMedia).

### Experimental Protocol

For ventricular pacing, a bipolar electrode was placed on the surface of the mid/base right ventricle. For sympathetic nerve stimulation (SNS), a 6F quadripolar catheter (2 mm electrode, 5 mm spacing) was inserted into the spinal cord to T1-T3 as previously described (20-22). As our previous studies demonstrated rabbit hearts have a strong response to SNS in a range of 5–15 Hz and 5–10 V for no longer than 60 s as measured by changes in HR (20), hearts in this study were stimulated with SNS at 8 Hz and 10 V for 60 s in sinus rhythm or for 13 s with ventricular pacing. Rate-matched baseline electrophysiological parameters were measured during continuous ventricular pacing at a pacing cycle length (PCL) of 200 ms with or without SNS. Incidence of arrhythmia was assessed following 60 s of SNS with ventricular pacing (progressively faster PCLs in 20 ms steps until loss of capture or induction of VT/VF). Responsiveness to β-adrenergic stimulation was assessed with isoproterenol (ISO; 30–100 nM) which was added to the perfusate.

### Optical Mapping Data Analysis

Data analysis was conducted using Optiq (Cairn Research) and Electromap (25) as described previously (10, 22, 26). Briefly, APD (APD_80_) and Ca^2+^ transient duration (CaTD_80_) were calculated as 80% repolarization minus activation time. APD dispersion was determined by dividing the range between the 5^th^ and 95^th^ percentiles of APDs within the field of view by the median APD value. The infarct region was defined as a signal to noise ratio (SNR) between 5–10 and an action potential (AP) rise time (Trise) >15 ms (27). The remote region was defined as an area of similar size but far from the infarct region and toward the base of the heart. The corresponding anatomical regions were also selected in sham hearts.

### Histology and Immunohistochemistry

Following optical mapping, 6 MI hearts were fixed in 4% paraformaldehyde and placed in 30% sucrose overnight. Short axis slices (approx. 2 mm) were cut (6 sections/heart), embedded in optimal cutting temperature (OCT) medium and frozen for storage at -80°C. OCT blocks were cryo-sectioned at 10 µm thickness and each section thaw-mounted onto positively charged slides (Acepix Biosciences Inc). Immunohistochemistry for CSPG and sympathetic nerve density with tyrosine hydroxylase (TH) was performed as previously described (7, 10, 28, 29). In brief, slides were rehydrated with phosphate buffered saline (PBS) and incubated in sodium borohydride (10 mg/mL; 3 x 3 min). Slides were then blocked with 2% bovine serum albumin (BSA, Sigma) in PBS containing 0.3% Triton X-100 (Sigma, PBS-T) for 1 hr and probed with primary antibodies targeting CSPG (1:300, Sigma CS-56) and TH (1:300, EMD Millipore) overnight. Secondary antibodies (Goat Anti-Mouse; Goat Anti-Rabbit respectively, both 1:500) conjugated to Alexa Fluor® 488 or to Alexa Fluor® 568 (Invitrogen) were applied to the samples for 90 mins. Glycerol in PBS (1:1) was used to mount coverslips. Sections were imaged on a Nikon Eclipse Ni microscope at 4x and 10x magnification with a FITC or TRITC filter (Ex 470/40 or 525/50 nm). A minimum of 5 images were taken of the infarct, border and remote zone. TH labelling was quantified with Nikon NIS-Elements Basic Research Microscope Imaging Software and the % tissue area that was TH+ was calculated. To identify the location and size of infarct regions, Masson’s trichrome staining (Trichrome Stain [Masson] Kit, Millipore-Sigma) was performed on adjacent sections as previously described (10).

### Statistics

Data are expressed as mean ± standard deviation (SD) for N animals. Statistical analysis was performed using GraphPad Prism 9. Normality was tested using the Shapiro-Wilk test and significance was assessed using student’s unpaired t*-*test, two-way ANOVA with repeated measures or mixed effects with Sidak’s multiple comparison correction, or Fisher’s exact test as appropriate and specified in the figure legend. Statistical significance was attained when *P*<0.05.

## RESULTS

### CSPGs and Sympathetic Nerve Density

Infarct presence and location were verified by Masson’s Trichrome staining (**Fig 1A**). TH labeling revealed visible sympathetic nerve fiber loss in the infarct vs. remote region (**Fig 1B**). CSPGs were present in the infarct and border zone of all (6/6) MI hearts examined (**Fig 1C**), with no CSPG+ signal observed outside of these areas. Sympathetic nerve density was quantified and found to be reduced in the infarct vs. remote region (**Fig 1D**; 0.56±0.40% vs. 6.41±2.78%, *P*=0.0005).

**Figure 1.**
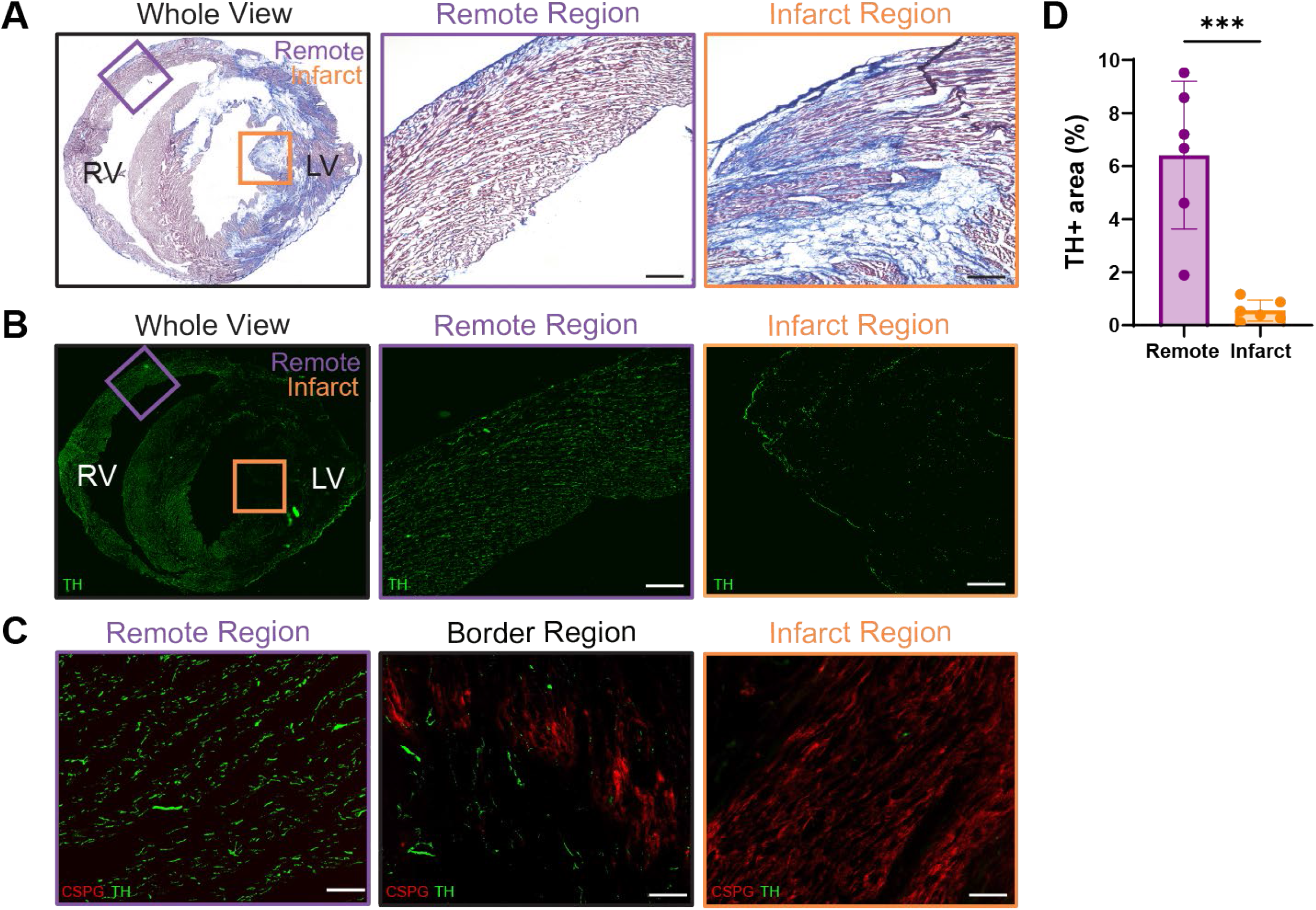
Masson’s trichrome, sympathetic nerve density, and CSPGs in MI hearts. **(A)** Masson’s trichrome staining showing fibrosis (blue) and myocardial tissue (red) in the whole view (left), remote (center), and infarct (right) regions. **(B)** Sympathetic nerve staining (TH, green) in the whole view (left), remote (center), and infarct (right) regions. **(C)** CSPG (red) and TH (green) in remote (left), border (center) and infarct (right) regions. **(D)** TH-positive tissue area. Data are mean±SD. N=6. ^* * *^*P*<0.001, by unpaired t-test. Scale bars = 100 µm.

### Effect of SNS on Heart Rate, Electrophysiology, and Arrhythmogenesis

The effect of SNS on electrophysiological parameters during sinus rhythm (**Fig 2**) and pacing (**Fig 3**) was assessed. During 60 s SNS in sinus rhythm, HR significantly increased by 10 s in both sham and MI and remained relatively stable for the remainder of stimulation (**Fig 2A**, *P=*0.02 and *P=*0.006, respectively). APD_80_ shortened with 60 s SNS in both sham and MI (*P*<0.001), with a trend for shorter APD in sham hearts, however no significant difference was observed between groups (**Fig 2B**, *P=*0.35). Interestingly, while APD_80_ dispersion remained constant during SNS in sham hearts, it gradually increased in MI hearts and peaked at 40 s where APD_80_ dispersion was increased vs. sham (**Fig 2C**, *P*=0.02).

**Figure 2.**
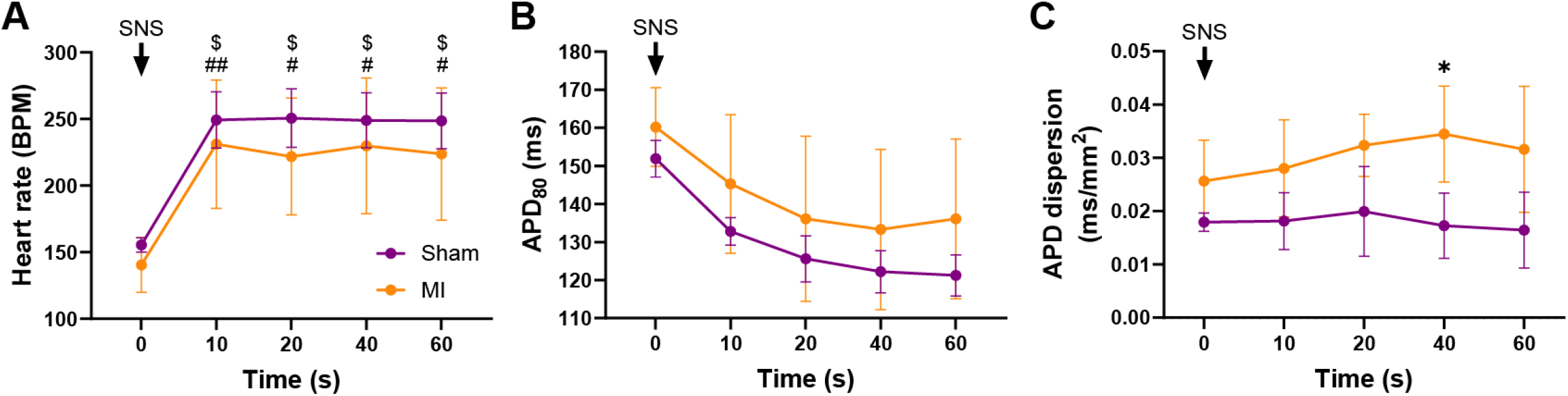
Effect of SNS on electrophysiological parameters during sinus rhythm. Changes in heart rate **(A)**, APD_80_ **(B)**, and APD dispersion **(C)** in response to SNS. Data are mean±SD. A: N=4-6/group (except 20s, N=4-5/group), B: N=3-5/group (except 60s, N=3-4/group), C: N=3-5/group. ^*^*P*<0.05 vs. sham, $/# *P*<0.05 vs. baseline in sham/MI, # # *P*<0.01 vs. baseline in MI, by two-way ANOVA with mixed-effects analysis (A, B), two-way ANOVA with repeated measures (C).

**Figure 3.**
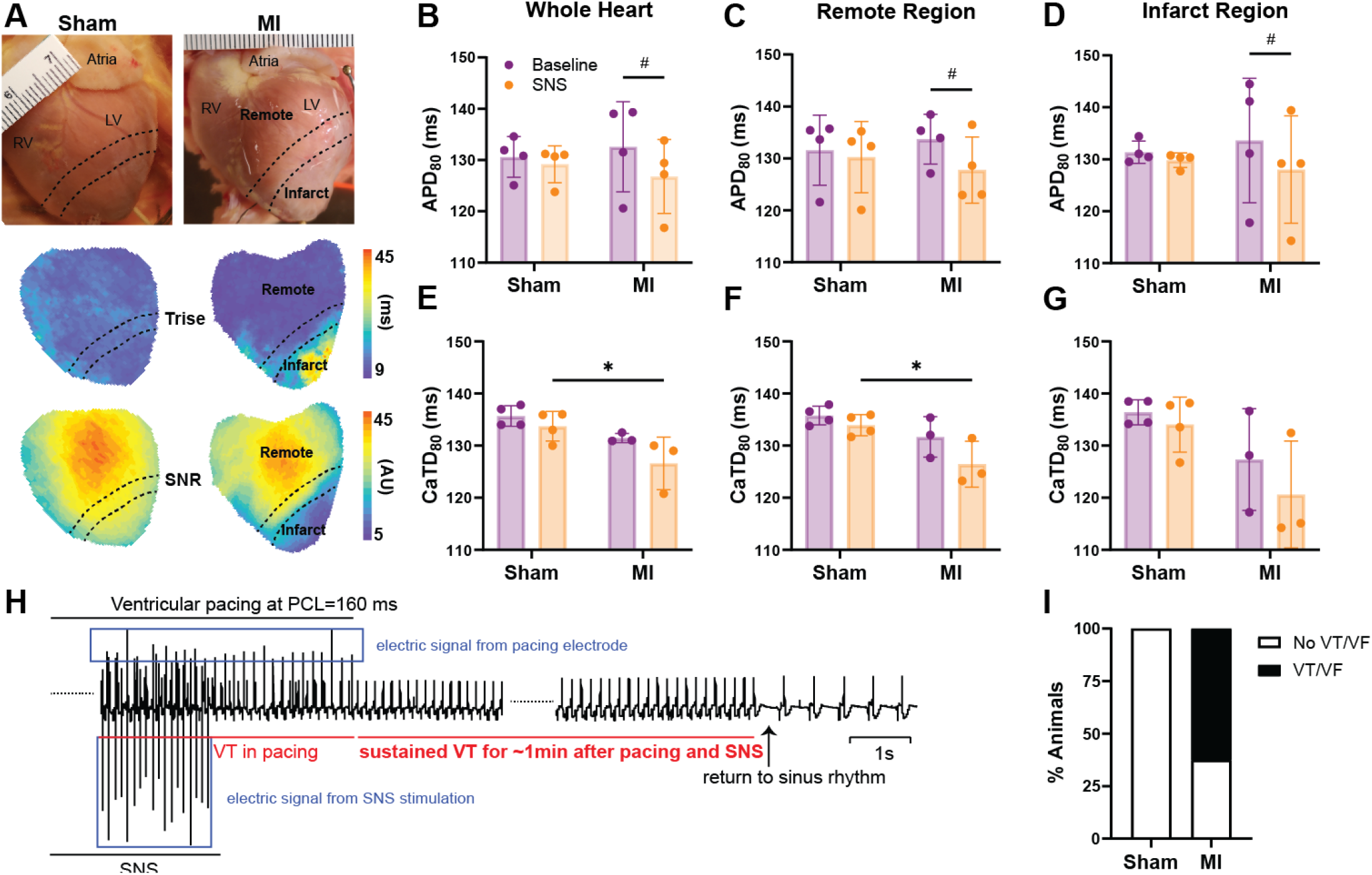
Effect of SNS on electrophysiological parameters and arrhythmia susceptibility during pacing. **(A)** Representative images and maps showing remote and infarct regions, with infarcts defined by CaT rise time (Trise) >15ms and signal to noise ratio (SNR) of 10-15. APD_80_ in the whole heart **(B)**, remote **(C)** and infarct **(D)** region at PCL=200 ms. CaTD_80_ in the whole heart **(E)**, remote **(F)** and infarct **(G)** region at PCL=200ms. **(H)** Example trace of ventricular tachycardia (VT). **(I)** Proportion of rabbits with VT or ventricular fibrillation (VF). Data are mean ± SD. B-D: N=4/group; E-G: N=3-4/group; I: N=4-8/group. ^*^*P*<0.05 MI vs. sham, #*P*<0.05 baseline vs. SNS in MI, by two-way ANOVA with repeated measures (B-G) and Fisher’s exact test (I).

To further determine the effect of SNS, rate-matched data (PCL=200ms) from whole heart, remote, and infarct regions with and without SNS were assessed. APD_80_ shortened in response to SNS in the whole heart, remote, and infarct regions of MI hearts, however no significant differences were observed in sham (**Fig 3B-D**). While no difference in CaTD_80_ with SNS was observed vs. baseline, SNS CaTD_80_ was shorter in MI vs. sham in whole heart and remote regions (**Fig 3E-F**, both *P*=0.02).

To assess susceptibility to arrhythmia, rapid ventricular pacing at increasing rates (shorter PCLs) was performed during SNS to induce VT/VF (**Fig 3H**). Though not significant, there was tendency for increased arrhythmias in MI with VT/VF induced in 5/8 MI hearts vs. 0/4 in sham (**Fig 3I**, *P*=0.08).

### β-adrenergic Responsiveness

As differences in electrophysiological responses to SNS may arise from altered sympathetic nerve density or function (e.g., local NE release), altered cardiomyocyte responsiveness to β-adrenergic stimulation, or a combination of both, we specifically evaluated β-adrenergic responsiveness across the ventricle using ISO. APD_80_ and CaTD_80_ shortened in sinus rhythm with ISO in both sham and MI (**Fig 4A**, both *P* = 0.0003 and **Fig 4C**, both *P* < 0.0001), with slightly greater relative ISO-mediated shortening of CaTD_80_ in MI vs. sham (**Fig 4D**, -42.60 ± 1.71% vs. -33.04 ± 4.67%, *P =* 0.02).

**Figure 4.**
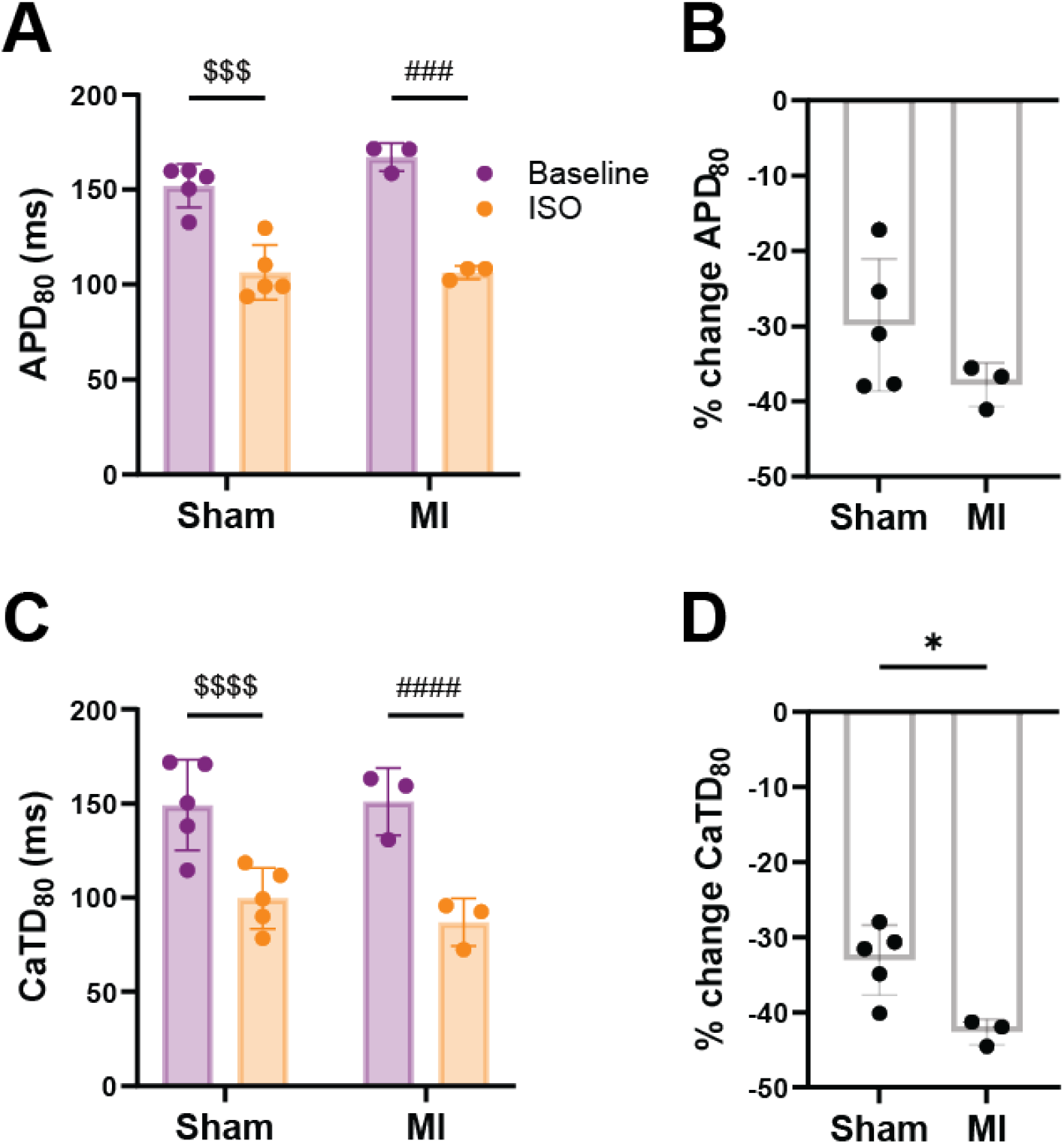
Effect of isoproterenol on APD_80_ and CaTD_80_ during sinus rhythm. APD_80_ **(A)**, % change in APD_80_ **(B)**, CaTD_80_ **(C)** and % change in CaTD_80_ **(D)** in response to isoproterenol (ISO) across the ventricle. Data are mean±SD. N=3-5/group. ^*^*P*<0.05 MI vs. sham, $$$/# # # *P*<0.001, $$$$/# # # # *P*<0.0001 baseline vs. ISO in sham/MI, by two-way ANOVA with repeated measures (A,C) or unpaired t-test (B,D).

## DISCUSSION

Here we show that: (1) CSPGs are present in the hypo-innervated infarct region of rabbit hearts with reperfused MI; (2) SNS increases APD dispersion and susceptibility for VT/VF in MI; and (3) despite significant hypo-innervation, MI hearts remain responsive to physiological nerve stimulation, potentially via preserved or slightly elevated β-adrenergic responsiveness.

Similar to previous studies in mice (7, 11), we found CSPGs present in both the infarct and border zone regions of post-MI rabbit hearts (**Fig. 1C**), with the infarct region having a significant reduction in sympathetic nerve density compared to the remote region (**Fig. 1B**). CSPGs, particularly when sulfated, mediate inhibition of nerve re-growth post-MI through binding to protein tyrosine phosphatase receptor σ (PTPRσ) present on sympathetic neurons. Deletion, modification, or pharmacological inhibition of PTPRσ, or CSPG sulfation have been shown to promote sympathetic reinnervation of the infarct (7, 11, 28, 30). Though sulfation of CSPGs was not assessed in this study, we found the presence of CSPGs to be associated with hypo-innervation, suggesting similar mechanisms of post-MI sympathetic remodeling between rodents and larger species. Interestingly, as CSPGs are absent in non-reperfused MI infarcts (e.g., chronic ligation models (11)), the cellular source of CSPGs and how CSPG expression is impacted by reperfusion remains an important area for future study.

Both sham and MI hearts demonstrated similar changes in HR and APD during SNS in sinus rhythm, however SNS resulted in a gradual increase in APD dispersion in MI but not sham hearts (**Fig. 2C**). These results are comparable to previous studies in models of MI and sympathetic denervation without MI (7, 10), in patients with post-infarct cardiomyopathy (9), and in a rabbit MI-induced heart failure model where APD restitution dispersion was increased (31). The increased APD dispersion we observed here is likely consequential of heterogeneous sympathetic innervation and potentially heterogeneous changes in β-adrenergic sensitivity post-MI. As increased APD dispersion or heterogeneity of refractoriness provides a substrate for unidirectional conduction block and reentry (13), this likely underlies the tendency for increased susceptibility to pacing-induced ventricular arrhythmias during SNS in post-MI hearts (**Fig. 3I**) (31).

Post-MI changes in responsiveness to SNS or β-adrenergic stimulation may also contribute to the observed differences in APD_80_ and CaTD_80_. Despite reduced sympathetic nerve density in the infarct region, APD_80_ shortened in response to SNS in MI (**Fig 3D**), indicating that the hypo-innervated rabbit myocardium is still sensitive and responsive to SNS, in line with a previous study (31). These findings could be due to changes in the amount of NE released from the remaining sympathetic nerves in the infarct region, changes in NE reuptake (which may result in increased NE diffusion from nearby nerves), or changes in β-adrenergic sensitivity of the myocardium. While ISO resulted in shortening of APD_80_ and CaTD_80_ from baseline values in both sham and MI hearts (**Fig 4A & 4C**), slightly greater relative changes in CaTD_80_ were observed in MI (**Fig 4D**), which may suggest enhanced responsiveness of Ca^2+^-handling proteins modulated by β-adrenergic activation (e.g. phospholamban phosphorylation on SERCA activity). Taken together, these results suggest that at 10±1 days post-MI, rabbit hearts are at least as sensitive to direct stimulation of β-adrenergic receptors as sham hearts, with perhaps slightly elevated sensitivity as indicated by the larger change in CaTD_80_. While β-adrenergic sensitivity likely changes throughout the post-MI time course, an exaggerated sympathetic effect has also been observed following 8 weeks of post MI-induced heart failure where intrinsic cardiac ganglia in the hilum region were enlarged (31).

### Conclusions

Here we used the innervated rabbit heart model to evaluate electrophysiological and sympathetic remodeling 10±1 days following reperfused MI. We found that CSPGs were present in the infarct region and were associated with significant sympathetic hypo-innervation, suggesting that the mechanisms of post-MI sympathetic remodeling may be similar in rodents and larger mammals. Moreover, despite hypo-innervation, post-MI hearts retained sympathetic responsiveness at this post-MI timepoint, while demonstrating pro-arrhythmic responses to SNS.

## DATA AVAILABILITY

Full datasets are available from the corresponding author upon reasonable request.

## AUTHOR CONTRIBUTIONS

A.G., G.A.N., B.A.H., Z.W., and C.M.R. conceived and designed research; A.G., C.E.R.S., L.W., J.L.C., S.T., S.D.S.F., B.W.M., and Z.W. performed experiments; A.G., C.E.R.S., L.W., J.L.C., B.W.M., G.A.N., B.A.H., Z.W., and C.M.R. analyzed data and interpreted results of experiments; A.G., C.E.R.S., Z.W., and C.M.R. prepared figures and drafted the manuscript; all authors edited and revised the manuscript and approved the final version of manuscript.

